# Xbra and Smad-1 response elements cooperate in PV.1 promoter to inhibit the early neurogenesis in *Xenopus* embryos

**DOI:** 10.1101/156992

**Authors:** Shiv Kumar, Zobia Umair, Jaeho Yoon, Unjoo Lee, SungChan Kim, Jae-Bong Park, Jae-Yong Lee, Jaebong Kim

**Affiliations:** Department of Biochemistry, Institute of Cell Differentiation and Aging, College of Medicine, Hallym University, Chuncheon, Gangwon-Do, 24252, Republic of Korea; Department of Electrical Engineering, Hallym University, Chuncheon, Gangwon-Do, 24252, Republic of Korea

**Keywords:** BMP-4/Smad-1 signaling,, FGF/Xbra signaling,, PV.1,, early neurogenesis,, *Xenopus.*

## Abstract

Crosstalk of signaling pathways plays crucial roles in cell fate determination, cell differentiation and proliferation. Both BMP-4/Smad-1 and FGF/Xbra signaling induce the expression of PV.1, leading to neural inhibition. However, BMP-4/Smad-1 and FGF/Xbra signaling crosstalk in the regulation of PV.1 transcription is still largely unknown. In this study, Smad-1 and Xbra physically interacted and regulated the PV.1 transcriptional activation in a synergistic manner. Xbra and Smad-1 directly bound within the proximal region of the PV.1 promoter and cooperatively enhanced the binding of an interacting partner within the promoter. Maximum cooperation was achieved in the presence of intact DNA binding sites of both Smad-1 and Xbra. Collectively, BMP-4/Smad-1 and FGF/Xbra signal crosstalk was required to activate the PV.1 transcription, synergistically. Suggesting that crosstalk of BMP-4 and FGF signaling facilitates the fine-tuning regulation of PV.1 transcription to inhibit neurogenesis during embryonic development of *Xenopus.*

**Summary statement:** FGF/Xbra positively regulates the PV.1 expression in the Xenopus via an unknown mechanism. Our study shows that both BMP-4/Smad-1 and FGF/Xbra exhibits a signaling crosstalk to regulate PV.1 transcription activation, promoting to ectoderm and mesoderm formation and inhibiting the early neurogenesis in *Xenopus.*

## Introduction

Bone morphogenetic protein-4 (BMP-4) signaling is actively involved in the germ-layer specification, dorsoventral axis formation, ventral mesodermal induction, neural inhibition, and cell differentiation in *Xenopus* (Hemmati-Brivanlou and Thomsen, 1995; Dale and Wardle, 1999). Blocking of BMP-4 signaling is facilitated by organizer-specific secreted BMP antagonists, including Chordin (Sasai et al., 1995; Piccolo et al., 1996), Noggin (Zimmerman et al., 1996), Follistatin (Fainsod et al., 1997), Cerberus (Bouwmeester et al., 1996) etc. Our previous study demonstrated that BMP-4/Smad-1 directly binds within endogenous PV.1 promoter region and induces the expression of PV.1, leading to the ventral mesodermal formation (Ault et al., 1996; Lee et al., 2011). Moreover, the C-terminal domain of PV.1 leads the inhibition of dorsal fate determination and neural inhibition (Hawley et al., 1995; Hwang et al., 2002; Hwang et al., 2003). Recent studies observed that FGF signaling plays a dual role to regulate the BMP-4/Smad-1 signaling (Pera et al., 2003; Yoon et al., 2014b). FGF signaling inhibits BMP-4/Smad-1 signaling through inhibiting the Smad-1 activity in an MAPK-dependent manner (Pera et al., 2003). However, FGF alone is not sufficient to induce neurogenesis in *Xenopus* embryos (Pera et al., 2003). Later on, a study showed that FGF signaling positively regulates expression of PV.1 (A transcriptional factor of BMP-4 signaling) in a Xbra-dependent manner, causing to neural inhibition (Yoon et al., 2014b). The involvement of BMP-4/Smad-1 and FGF-Xbra signaling may facilitate a signaling crosstalk in the regulation of PV.1 expression during embryonic development of *Xenopus.* (Lee et al., 2011; Yoon et al., 2014b). However, the detailed molecular mechanism of BMP-4/Smad-1 and FGF/Xbra-mediated signaling crosstalk in the regulation of PV.1 transcription is largely unknown.

PV.1 is one of the homeobox-containing Xvent family members (Ault et al., 1996). Xvent family is divided into two subfamilies (Xvent-1 and Xvent-2 subfamilies) on account of amino acids differences. Xvent-1 subfamily; PV.1, Xvent-1, Xvent-1B, and PV.1A (Gawantka et al., 1995; Ault et al., 1996; Rastegar et al., 1999) while Xvent-2 subfamily; Xvent-2, Xbr-1b, Xom, Vax and Xvent-2B (Ladher et al., 1996; Onichtchouk et al., 1996; Papalopulu and Kintner, 1996; Schmidt et al., 1996; Rastegar et al., 1999). PV.1 is a direct target of BMP-4/Smad-1 signaling and vitally implicated in ectoderm and ventral mesodermal formation. The proximal promoter region of PV.1 contains various cis-acting response elements for transcriptional activation of BMP-4 downstream targets, including Xvent-2, Oaz, and Smad-1 (Lee et al., 2011). Moreover, PV.1 negatively regulates expression of neurogenesis-promoting genes, including *Chordin, Zic3, FoxD5b, Ngnr,* and *N-CAM* (Melby et al., 1999; Trindade et al., 1999; Lee et al., 2002; Yoon et al., 2014b). However, the detailed molecular mechanism of PV.1-mediated inhibition of early neurogenesis is still awaited to delineate.

FGF/Xbra signaling is well-known to promote mesoderm formation, cell fate determination, and anterior-posterior (A-P) patterning of neural tissue. FGF signaling causes the mesoderm formation through the activation of the autocatalytic loop (FGF/Ras/Xbra/AP-1) in *Xenopus* embryos (Kim et al., 1998; Gamse and Sive, 2000; Weisinger et al., 2008). Xbra is one of the well-known members of a T-box gene family that acts as a transcriptional activator through its C-terminal domain. Ectopic expression of Xbra facilitates posterior mesoderm and notochord formation during vertebrate development (Saka et al., 2000). Moreover, Xbra also stimulates ventral and lateral mesoderm formation in animal cap explants. A study documented that C-terminus truncated mutant of Xbra can trigger the early neurogenesis without showing Xbra-mediated mesodermal activity in ectodermal animal cap explants, causing the anterior neural formation (Rao, 1994). Recent studies have been documented that T-box transcriptional activator proteins bind to a consensus sequences (A/G)(A/T)(A/T)NTN(A/G)CAC(C/T)T within promoter region of its targeted genes, promoting to transcription activation of targeted genes (Conlon et al., 2001; Kusch et al., 2002). Our previous study addressed that FGF signaling promotes PV.1 expression, causing to neural inhibition while dominant-negative Xbra (DN-Xbra) can trigger early neurogenesis in ectoderm through inhibiting the PV.1 expression (Yoon et al., 2014b). This study suggests that FGF signaling induces the PV.1 expression in a Xbra-dependent manner. Moreover, Xbra may regulate the transcriptional activation of PV.1. However, the detailed mechanism of Xbra-mediated transcriptional activation of PV.1 expression is not completely understood.

A previous study reported that C-terminal phosphorylated Smad-1 physically interacts with N-terminal of Xbra during early *Xenopus* development (Messenger et al., 2005). However, the regulatory function of Xbra-Smad-1 interaction is still uncovered. Our present study demonstrated that Xbra directly bound on Xbra response elements, ATCACACTT (XbRE, within −70bp~-62bp) and positively induced the PV.1 transcription. Results showed that Xbra and Smad-1 cooperated synergistically to regulate the PV.1 transcription during *Xenopus* development. Moreover, ChIP assay indicated that Smad-1 and Xbra directly bound to their respective consensus sequences within the proximal region of the endogenous PV.1 promoter and enhanced the binding of their respective interacting partners to the promoter region. Additionally, we observed that Xbra and Smad-1 interaction showed target specificity in the Vent family because Xbra and Smad-1 were not able to induce Xvent2 expression, synergistically. Taken together, evidence concluded that FGF/Xbra and BMP-4/Smad-1 signaling positively involves in transcriptional activation of PV.1 and cooperates synergistically to induce the PV.1 expression, leading to dorsoventral patterning and ventral mesodermal formation. Moreover, BMP-4/Smad-1 and FGF/Xbra signaling demonstrate a crosstalk in the inhibition early neurogenesis and PV.1 transcriptional activation during early *Xenopus* development.

## Results

### 1. Xbra negatively regulates the neurogenesis in early *Xenopus* development

Our previous study demonstrated that FGF/Xbra induces expression of BMP downstream target gene, *PV.1* which negatively regulates early neurogenesis in ectoderm in *Xenopus* (Yoon et al., 2014b). Thus, we asked whether Xbra induces PV.1 expression in BMP-4 inhibition condition, causing to neural inhibition. RT-PCR results showed that co-injection of Xbra with DNBR induced expression of PV. 1 and suppressed the expression of the early neural gene, *FoxD5a,* at early gastrula stage (Figure 1A). The result suggested that Xbra positively regulated the PV.1 expression in BMP inhibition condition to inhibit the neurogenesis. Moreover, Xbra co-injection strongly inhibited the expression of late neural genes, including *N-CAM, Ngnr,* and *Otx20* which were highly expressed by DNBR alone at tailbud stage (Figure 1B). These results suggested that Xbra induced PV.1 expression in BMP-4 inhibited condition and negatively regulated the early neurogenesis during embryonic development of *Xenopus*. This finding supports to our previous study that FGF/Xbra-mediated induction of PV.1 negatively regulates the early neurogenesis in *Xenopus* embryos (Yoon et al., 2014b).

**Figure 1.**
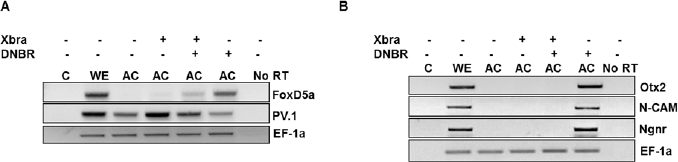
Ectopic expression of Xbra negatively regulates the neurogenesis in early *Xenopus* development. Xbra (1ng) mRNA was injected with or without DNBR (1ng) mRNA at one cell stage to dissect the animal cap at stage 8 and harvested at stage 11, and 24. The relative gene expression was analyzed by RT-PCR. (A) Xbra increases *PV.1* expression and reduces *FoxD5a* expression while DNBR reduces PV.1 expression and induces FoxD5a expression. All experiments performed at stage 11. (B) Xbra injection reduces the expression of *N-CAM, Ngnr*(Pan-neural markers), and *Otx2* (Anterior neural marker) and while DNBR induces the expression of *N-CAM, Ngnr,* and *Otx2.* All experiments performed at tailbud stage 24.

### 2. Inhibition of neurogenesis by Xbra is mediated through PV.1

Our previous study documented that BMP-4/Smad-1 directly binds to the endogenous promoter of PV.1 and induces PV.1 expression during *Xenopus* development (Lee et al., 2011). Our recent study showed that FGF/Xbra induces PV.1 expression and negatively regulates the early neurogenesis (Yoon et al., 2014b). However, it is still questionable whether Xbra-mediated inhibition of neurogenesis was dependent on PV.1. We co-injected PV.1 MOs with Xbra and DNBR to perform RT-PCR at early gastrula and tail-bud stage. Knockdown of PV.1 by PV.1 MOs strongly augmented and recovered the expression of early and late neural genes, including *FoxD5a, N-CAM, Ngnr,* and *Otx2* which were reduced by Xbra. PV.1 MOs specifically reduced *PV.1* expression in both condition absence and presence of Xbra (Figure 2A and 2B). These results showed that Xbra facilitated the inhibition of neurogenesis in a PV.1-dependent manner. Taken together, the results suggested that 5’-flanking region of PV.1 may probably contain the Xbra response elements (XbRE) within the PV.1 promoter region. Moreover, it also suggested that PV.1 is a key molecule which made a bridge and established a signaling crosstalk between BMP-4/Smad-1 and FGF/Xbra to inhibit the early neurogenesis in *Xenopus* embryos.

**Figure 2.**
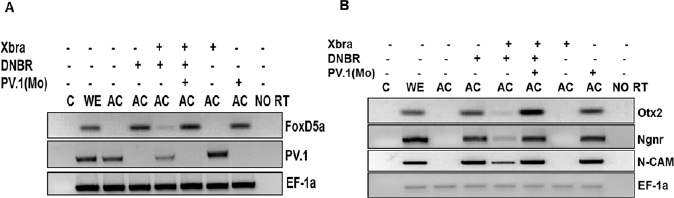
Inhibition of neurogenesis by Xbra mediates through PV.1. PV.1 MOs (1ng) were injected with or without DNBR and Xbra mRNA at one cell stage and dissected the animal cap at stage 8. The dissected animal caps were harvest until stage 11 and 24 in 0.5X L-15 medium. The relative gene expression was analyzed by RT-PCR. (A and B) PV.1 MOs reduces the PV.1 expression and increases *FoxD5a* expression while PV.1 MOs augments expression of *N-CAM, Ngnr,* and *Otx2* at tailbud stage 24.

### 3. Xbra response element (XbRE) is identified in the PV.1 promoter

5’-flanking region of PV.1 contains various cis-acting response elements, including Xvent-2, Oaz and Smads and triggers the PV.1 transcriptional activation for regulating BMP-4 signaling (Lee et al., 2011). Our above findings suggested that PV.1 promoter region may contain putative cis-acting XbRE. To evaluate the presence of cis-acting XbRE within PV.1 promoter region, we injected PV.1 (−2525) promoter reporter construct with or without DN-Xbra, and Xbra, separately. The results showed that DN-Xbra decreased the relative promoter activity of PV. 1 (-2525) up to 2.5-fold while Xbra mRNA increased the relative promoter activity of PV.1 (−2525) up to 3.5-fold as compared to PV.1 (−2525) alone (Figure 3A and 3B). Results evidenced that Xbra positively regulated PV.1 transcription activation. Moreover, PV.1 promoter region contained putative cis-acting XbRE. Therefore, we generated different serially deleted PV.1 promoter constructs to map out the cis-acting XbRE within PV.1 promoter region (Figure 3C). We co-injected different serially deleted PV.1 promoter constructs with and without Xbra mRNA. Results showed that Xbra increased the relative promoter activity of serially deleted PV.1 promoter constructs up to 1.5-4.0-fold as compared to PV.1 promoter constructs alone, separately (Figure 3D). These findings strongly suggested that PV.1 (−103) promoter construct contained cis-acting XbRE to trigger PV.1 transcription. We further tested PV.1 (−103) promoter construct with and without Xbra mRNA in a dose-dependent manner to follow the relative promoter activity of PV. 1. The result showed that Xbra increased the relative promoter activity of PV.1 (−103) up to 1.5-3.0-fold in a dose-dependent manner (Figure 3E). These results strongly provide evidence that PV.1 (−103) promoter construct contained cis-acting XbRE and induced the PV. 1 transcription.

**Figure 3:**
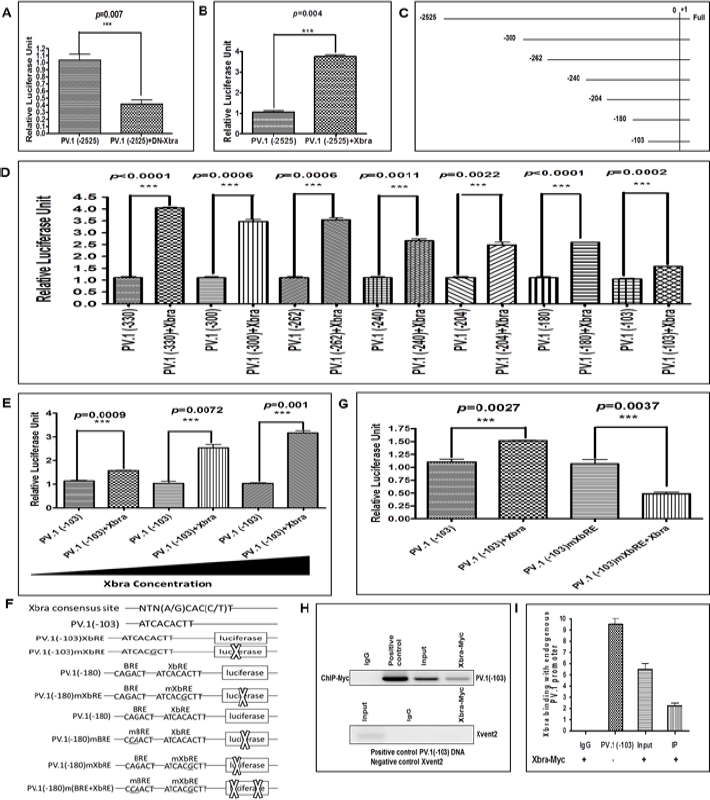
Identification of Xbra response elements (XbRE) within PV.1 promoter region. Different serially deleted PV.1 (40pg) promoter construct was co-injected with or without 1ng either DN-Xbra or Xbra at 1-cell stage and grown till stage 11 in 30% MMR to measure relative promoter activity. (A) DN-Xbra decreases the relative promoter activity of PV.1 (−2525) promoter region as compared to PV.1 (−2525) promoter region alone. (B) Xbra increases the relative promoter activity of PV.1 (−2525) promoter region as compared to PV.1 (−2525) alone. (C and D) Xbra increases the relative promoter activity of serially deleted promoter construct of PV.1. (E) Xbra increases the relative promoter activity of PV.1 (−103) promoter construct in a dose-dependent manner. (F) The loss-of-function study was performed with PV.1 (−103 and −180) promoter construct at Xbra and Smad-1 binding consensus sequences. (G) Xbra-mediated induction in relative promoter activity of PV.1 (−103) is abolished with mutated PV.1 (−103)mXbRE. (H and I) In ChIP assay, Xbra-Myc (1ng) was injected at 1-cell stage and grown till stage 11 in 30% MMR. Anti-Myc antibody was used to immunoprecipitate the endogenous PV.1 promoter region. All binding was measured by PCR method with specific primers. PV.1 (-103) promoter DNA used as positive control and Xvent2 coding region used as a negative control for all ChIP experiments All relative promoter activity data are shown as mean ± SE.

Previous studies have been addressed the putative T-box transcription factors binding consensus sequences NTN(A/G)CAC(C/T)T within the promoter region of targeted genes (Conlon et al., 2001; Kusch et al., 2002). Thus, we mapped out PV.1 (−103) promoter nucleotide sequence and found putative Xbra binding consensus sequences, ATCACACTT (XbRE, within −70bp~−62bp) upstream of putative transcription initiation site of PV.1. Thus, we mutated one nucleotide (ATCACACTT-ATCACGCTT) within putative XbRE containing PV.1 (−103 and −180) promoter constructs at −65bp (Figure 3F). PV.1 (−103) and mutated PV.1 (−103)mXbRE promoter constructs were tested with and without Xbra and measured the relative promoter activity with and without Xbra. Notably, Xbra-mediated transcriptional activation of PV.1 was abolished in PV.1 (−103)mXbRE as compared to PV.1 (−103) (Figure 3G). This finding firmly evidenced that PV.1 promoter constructs encompassed cis-acting XbRE within the proximal (−70bp~−62bp) region of the PV. 1 promoter. We further asked whether Xbra directly binds within the promoter region of PV.1 to activate the PV.1 transcription. ChIP assay showed that Xbra directly bound within the proximal region of the endogenous PV.1 promoter (Figure 3H and I). These findings collectively concluded that PV. 1 was one of the direct targets of Xbra. In addition, PV. 1 required direct binding of Xbra within proximal promoter region for triggering its transcription activation.

### 4. Xbra and Smad-1 synergistically regulate the PV.1 transcription

Previously, a study demonstrated that Smad-1 is a co-transcriptional activator of BMP signaling and directly binds with cis-acting BMP-4 response elements (BRE; CAGACT, −180bp to −162bp) within upstream region of putative transcriptional initiation site of PV.1 (Lee et al., 2011). Moreover, another study has been documented that phosphorylated C-terminal domain of Smad-1 physically interacts with the N-terminal domain of Xbra. Furthermore, this study suggested that Smad-1-Xbra interaction may induce the expression of ventral genes (Messenger et al., 2005). It has been evidenced that Xbra and Smad-1 both positively regulated the PV.1 expression and their cis-acting elements existed in close vicinity within the proximal region of the PV.1 promoter. However, the regulatory function of Smad-1-Xbra interaction and existence of their response elements in close vicinity is still unknown.

Thus, we further postulated that whether Xbra and Smad-1 may cooperate synergistically to activate PV.1 transcription during embryonic development of *Xenopus.* PV.1 (−103) promoter construct co-injected with both Xbra, and Smad-1 to measure relative promoter activity of PV.1. Results showed that concomitant overexpression of Smad-1 and Xbra increased up to 3.5-fold relative promoter activity of PV.1 (−103) while Xbra alone increased only 1.5-fold relative promoter activity of PV.1 (−103) as compared to PV.1 (−103) alone (Figure 4A). The result indicated that Smad-1 cooperated synergistically with Xbra to activate PV.1 transcription without binding to its cis-acting element because PV.1 (−103) promoter construct did not include the cis-acting element for Smad-1 (BRE). We then asked how much fold Xbra increases relative promoter activity of PV.1 in a synergistic manner with Smad-1 when PV.1 promoter included both consensus cis-acting BRE and XbRE. We co-injected PV.1 (−180) promoter construct, which contained both consensus cis-acting elements BRE and XbRE, with Xbra and Smad-1, either in combination or separately. The results showed that concomitant overexpression of Smad-1 and Xbra robustly increased relative promoter activity of PV.1 (−180) up to 15-fold while Xbra co-injection increased 2.5-fold relative promoter activity of PV.1 (−180) while Smad-1 increased 3.5-fold promoter activity PV.1 (−180) as compared to PV.1 (−180) alone (Figure 4B). The results strongly indicated that both Smad-1 and Xbra cooperated synergistically to activate the PV.1 transcription. Moreover, these results collectively suggested that Smad-1 and Xbra-mediated synergistic activation of PV.1 transcription established a signaling crosstalk between FGF/Xbra and BMP-4/Smad-1 signaling. To evaluate the Xbra and Smad-1-mediated synergistic regulation of PV.1 transcription, we injected PV.1 (−103) and mutated PV.1 (−103)mXbRE promoter constructs with both Xbra and Smad-1, in combination and separately. This result showed that Xbra and Smad-1-mediated synergistic stimulation of PV.1 (−103) promoter construct was completely abolished with PV.1 (−103)mXbRE (Figure 4C, bar 4-6) as compared to PV.1 (−103)-Xbra (Figure 4C, bar 2). The result indicated that XbRE played a significant role in a Xbra-mediated regulation of PV.1 transcriptional activation. Moreover, XbRE actively participated in the BMP-4/Smad-1 and FGF/Xbra-mediated signaling crosstalk to regulate the synergistic activation of PV.1 transcription.

**Figure 4:**
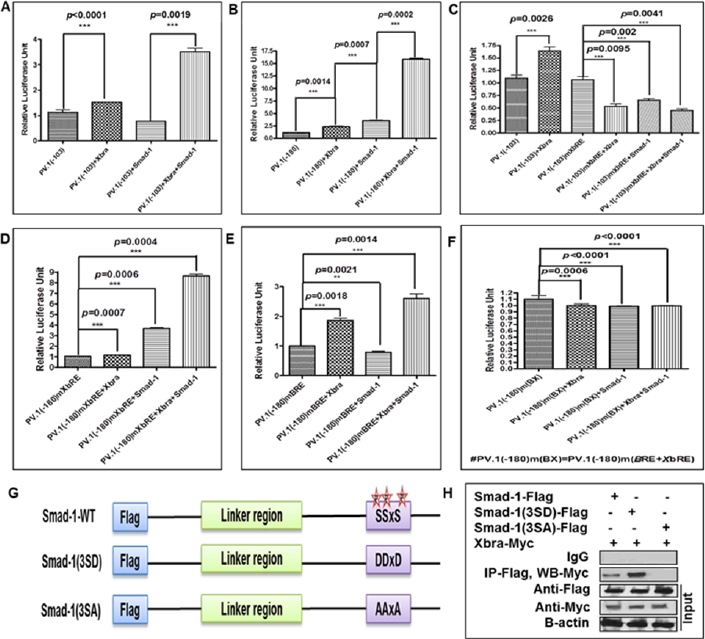
Xbra and smad-1 synergistically regulate the PV.1 transcription. Different PV.1 promoter constructs were co-injected with or without Smad-1, and Xbra mRNA, either in a combination or separately, at 1-cell stage and grown till stage 11 in 30 *%* MMR. Relative promoter activity was measured at stage 11. (A and B) The co-injection of Xbra-Smad-1 strongly increases the relative promoter activity of PV.1 (−103 and −180) as compared to PV.1 (−103 and −180)-Xbra. (C) The co-injection of Xbra-Smad-1 is not able to stimulate relative promoter activity of PV.1 (−103)mXbRE as compared to Figure 4A. (D) The relative promoter activity of PV.1 (−180)mXbRE promoter construct significantly decreases in Xbra-Smad-1 injected embryos as compared to Figure 4B. (E) The concomitant overexpression of Smad-1 and Xbra robustly decreases the relative promoter activity of PV.1 (−180)mBRE as compared to Figure 4B. (F) Xbra and Smad-1-mediated induction in relative promoter activity of PV.1 (−180) completely diminish with doubly mutated PV.1 (−180)m(BRE+XbRE) as compared to Figure 4B, 4D, and 4E. (G and H) 1ng of Smad-1-Flag, phospho-mimic Smad-1 3SD-Flag, and phosphorylation dead-mutant Smad-1 3SA-Flag were co-injected with Xbra-Myc (1ng) at 1-cell stage and collected the total protein at stage 11 to perform the immuno-precipitation with anti-
Flag antibody. All relative promoter activity experiments performed in triplicate. All relative promoter activity data are shown as mean ± SE.

Concomitant overexpression of Xbra and Smad-1 increased the relative promoter activity of PV.1 (−103 and −180) up to 3.5-fold and 15-fold as compared to PV.1 (−103) and PV.1 (−180) alone, respectively (Figure 4A and 4B, bar 4). We thus postulated that whether both cis-acting elements of BMP-4/Smad-1 response element (BRE) and FGF/Xbra response element (XbRE) actively participate in synergistic activation of PV.1 transcription.

To compare the contribution of each response element (BRE or XbRE) in synergistic activation of PV.1 transcription, we mutated BRE and XbRE within different PV.1 (−180) promoter constructs, separately and generated PV.1 (−180)mXbRE and PV.1 (−180)mBRE promoter constructs. The PV.1 (−180)mXbRE promoter construct was co-injected with and without Smad-1, and Xbra, in combination or separately. As expected, the result showed that Xbra-mediated stimulation was completely abolished (Figure 4D, 2nd bar) and Smad-1 mediated stimulation was sustained with a 3.5-fold increase in the presence of Smad-1 (Figure 4D, 3rd bar). Interestingly, site-directed mutagenesis of XbRE within PV.1 (−180) promoter construct, PV.1 (−180)mXbRE, still contained synergistic activation when Xbra was co-injected with Smad-1 (Figure 4D, 4th bar). Although, Xbra and Smad-1-mediated relative promoter activity of PV.1 (−180)mXbRE was decreased up to 8-fold compared to 15-fold which increased by Xbra and Smad-1 co-injected embryos with wild-type of PV.1 (−180) promoter construct (Figure 4B, 4th bar). The result indicated that XbRE played a significant role in synergistic activation of PV.1 transcription with Smad-1 and Xbra. To delineate the role of BRE in Smad-1 and Xbra-mediated synergistic activation of PV.1 transcription, we examined BRE mutated PV.1 promoter (PV.1 (−180)mBRE with and without Smad-1, and Xbra, in combination or separately. The results showed that Smad-1-mediated stimulation was completely abolished (Figure 4E, 3rd bar), but Xbra-mediated stimulation was sustained with the 1.5-fold increase in the presence of Xbra (Figure 4E, 2nd bar). Site-directed mutagenesis of BMP-4/Smad-1 response element (BRE) within PV.1 (−180) promoter construct (PV.1 (−180)mBRE) still sustained some synergistic activation when Smad-1 was co-injected with Xbra (Figure 4E, 4th bar). While, the relative promoter activity of PV.1 (−180)mBRE was dramatically decreased up to 2.5-fold as compared to 15-fold increase with the wild type of PV.1 (−180) promoter construct in the presence of both Xbra and Smad-1 (Figure 4B, 4th bar). Moreover, concomitant overexpression of Xbra and Smad-1 increased relative promoter activity of PV.1 (−180)mXbRE promoter construct up to 8.5-fold (Figure 4D, 4th bar).

To examine the role of both cis-acting elements (BRE and XbRE) in PV.1 (−180) promoter construct, we mutated both cis-acting consensus XbRE and BRE within PV.1 (−180) promoter construct as shown in Figure 3F. We co-injected doubly mutated PV.1 (−180)m(BRE+XbRE) promoter construct with Smad-1, and Xbra, in combination or separately. The results showed that Xbra and Smad-1-mediated stimulation in relative promoter activity of PV.1 (−180) promoter construct was completely abolished in PV.1 (−180)m(BRE+XbRE) (Figure 4F) while concomitant overexpression of Smad-1 and Xbra increased up to 16-fold, 8.5-fold and 2.5-fold relative promoter activity with wild-type PV.1 (−180), PV.1 (−180)mXbRE and PV.1 (−180)mBRE, respectively (Figure 4B, 4D and 4E, bar 4). The results indicated that both consensus cis-acting BRE and XbRE were required for maximum activation of PV.1 transcription in synergistic manner. Moreover, BMP-4/Smad-1 and FGF/Xbra signaling crosstalk was played a critical role in a PV. 1-dependent neural inhibition during early *Xenopus* development. We thus postulated whether Xbra and Smad-1 physically interact to regulate the synergistic activation of PV.1. We co-injected Xbra with the different construct of Smad-1; Smad-1 (wild-type), C-terminal phospho-mimic Smad-1 (3SD), and C-terminal phospho-dead mutant Smad-1 (3SA) and harvested injected embryos till stage 11 to perform immunoprecipitation assay (Figure 4G). The results showed that Xbra physically interacted with Smad-1 (wild-type) and phospho-mimic Smad-1 (3SD) while the interaction of Xbra with phospho-dead mutant Smad-1 (3SA) was not detected (Figure 4H). This result suggested that C-terminal phosphorylation of Smad-1 played a significant role in Xbra-Smad-1 interaction and notably required for synergistic regulation of PV.1 transcription activation. Moreover, the results showed that Smad-1-Xbra interaction was critically required for establishment of BMP-4/Smad-1 and FGF/Xbra-mediated signaling crosstalk which regulated the transcriptional activation of PV.1. This result supported to the previous study that Smad-1 C-terminal phosphorylation is remarkably required for Smad-1-Xbra interaction for stimulating the expression of the ventral genes (Messenger et al., 2005).

### 5. Xbra strongly enhances the DNA binding of Smad-1 on the endogenous PV.1 promoter region

For further confirmation of Smad-1 and Xbra-mediated synergistic regulation of PV.1 transcription, we assumed whether both Smad-1 and Xbra stimulates DNA binding of their interacting partner within the endogenous PV.1 promoter region. We co-injected Myc-Xbra mRNA with and without Smad-1 to perform the ChIP-PCR assay with anti-Myc antibody (Blythe et al., 2009). The result showed that ectopic expression of Smad-1 stimulated Xbra binding within the proximal region of the endogenous PV.1 promoter (Figure 5A and B). Thus, we further asked whether Xbra stimulates Smad-1 binding within endogenous PV.1 promoter region and restores the inhibition of Smad-1, mediated by FGF/MAPK signaling (Schier, 2001; Pera et al., 2003). We co-injected Flag-Smad-1 mRNA with and without Xbra and performed ChIP-PCR assay with anti-Flag antibody. Surprisingly, this result showed that Xbra robustly enhanced the Smad-1 binding within the proximal region of endogenous PV.1 promoter (−180bp to −162bp) upstream of putative transcription initiation site (Figure 5C and D). Further, we postulated whether the physical interaction of Smad-1 and Xbra regulates expression of other gene of the Vent family, we tested the Xvent promoter region with Smad-1 and Xbra both in separately, and in combination. Results exhibited that Smad-1 and Xbra increased relative promoter activity of Xvent2 (−1031) promoter construct and induced the expression of Xvent2, separately, but the concomitant injection of Smad-1 and Xbra was not able to regulate the expression of Xvent2 in a synergistic manner (Supplementary Figure 1). These results collectively indicated that both Xbra and Smad-1 directly bound within the proximal promoter region of PV.1. However, the direct binding of Xbra and Smad-1 within PV.1 promoter region was not critically required for synergistic regulation of PV.1 transcription. These results strongly suggested that Smad-1 played a significant role in a synergistic regulation of PV.1 transcription and also in BMP-4/Smad-1 and FGF/Xbra-mediated signaling crosstalk. Moreover, Xbra not only positively regulated the PV.1 transcription activation but also strongly stimulated the BMP-4/Smad-1 signaling. Also, the result suggests that synergistic effect of Smad-1 and Xbra may restore the activity of FGF/MAPK-mediated inhibition of BMP-4/Smad-1 signaling. Furthermore, suggested that PV. 1 may be a novel target of Smad-1 and Xbra interaction in a Vent family.

**Figure 5.**
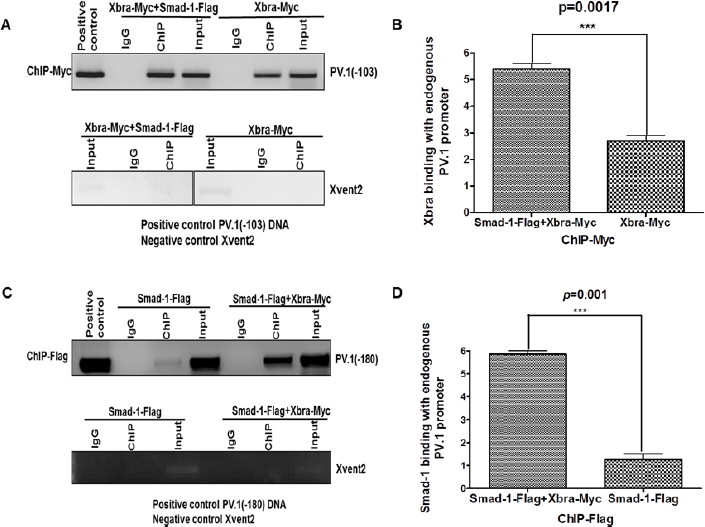
Physical binding of Smadl and Xbra stimulates promoter activity of PV.1 and is dependent on DNA binding, but not absolutely require DNA binding for cooperative stimulation. In chromatin immunoprecipitation assay, Xbra-Myc (1ng) was injected with or without Smad-1 at 1-cell stage and harvested the embryos till stage 11 in 30% MMR. (A and B) ChIP assay is performed with anti-Myc antibody and found that Smad-1 increases the Xbra binding with endogenous PV1 promoter region as compared to Xbra alone. (C and D) We also ChIP assay is performed with anti-Flag antibody and observed that Xbra also increases the Smad-1 binding with endogenous PV.1 promoter region as compared to Smad-1 alone. All binding was measured by PCR method with specific primers. PV1 (−103 and −180) promoter DNA used as positive control while Xvent2 coding region primers used for PCR as a negative control for all ChIP experiments.

## Discussion

Studies demonstrated that both BMP-4/Smad-1 and FGF/Xbra signaling positively regulates the transcription of the BMP-4 downstream target gene, *PV.1* to promote embryos ventralization and ventral mesodermal formation (Lee et al., 2011; Yoon et al., 2014b). These studies established a signaling crosstalk between BMP-4/Smad-1 and FGF/Xbra signaling. However, the significance and mechanism of this signaling crosstalk are still mostly awaited to delineate. BMP-4/Smad-1 facilitates the ectoderm formation and inhibits the neural induction during early *Xenopus* development. Inhibition of BMP signaling is required for inducing the early neurogenesis in *Xenopus* development (Hawley et al., 1995; Wilson and Hemmati-Brivanlou, 1995; Xu et al., 1995; Hemmati-Brivanlou and Melton, 1997). Furthermore, FGF/Xbra signaling actively participates in the lateral mesodermal formation and A-P patterning of neural tissue (Kim et al., 1998; Gamse and Sive, 2000; Weisinger et al., 2008). Previous studies demonstrated that FGF/Xbra signaling catalyzes the inhibitory phosphorylation of Smad-1 at linker region and inhibits the BMP-4/Smad-1 signaling (Schier, 2001; Pera et al., 2003). Surprisingly, FGF/Xbra-mediated inhibition of BMP-4 signaling alone is not sufficient to trigger the neurogenesis in ectodermal explant (Wilson et al., 2001; Delaune et al., 2005). A recent study documented that FGF/Xbra signaling also stimulates the PV.1 expression and inhibits the ectoderm-neuroectoderm transition in ectodermal animal cap explant of *Xenopus* embryos (Yoon et al., 2014b). However, the detailed mechanism of FGF/Xbra-mediated regulation of PV.1 activation was not delineated. BMP-4/Smad-1 and FGF/Xbra both positively regulates the PV.1 expression during early development of *Xenopus* embryos (Lee et al., 2011; Yoon et al., 2014b). The molecular mechanism of BMP-4/Smad-1 and FGF/Xbra-mediated signaling crosstalk in context to the positive and synergistic regulation of PV.1 expression is still largely unknown. Moreover, Xbra-mediated neural inhibition is still awaited to delineate. In this study, we outlined the molecular mechanism of FGF/Xbra and BMP-4/Smad-1-mediated signaling crosstalk and synergistic activation of PV.1 transcription to inhibit early neurogenesis in *Xenopus* development.

### Xbra inhibits the neurogenesis in a PV.1 dependent manner during *Xenopus* development

Our previous study provides evidence that ectopic expression of Xbra induces the PV.1 expression and inhibits the early neurogenesis (Yoon et al., 2014b). However, whether Xbra induces the PV.1 expression in the absence of BMP-4 signaling and inhibits the early neurogenesis was largely unknown. Xbra induces the expression of ventral gene *PV.1* in the absence of BMP-4 signaling (BMP-4 is inhibited by DNBR) and inhibits early neurogenesis through downregulating the expression of early and late neural markers genes, including

*FoxD5a, Ngnr, N-CAM* and *Otx2,* and (Figure 1A and 1B). This finding supports our previous study that Xbra induces the PV.1 expression and inhibits the early neurogenesis (Yoon et al., 2014b). However, this is still questionable that whether Xbra-mediated neural inhibition is facilitated in a PV.1-dependent manner. We demonstrated that PV.1 MOs-mediated knockdown of PV.1 restores the Xbra-mediated inhibition of early neurogenesis in ectodermal animal cap explants. Moreover, the ectopic expression of PV.1 MOs increases the expression of early and late neural marker genes, including *FoxD5a, Ngnr, N-CAM,* and *Otx2,* which decreased by Xbra, and PV.1 MOs also decreases the Xbra-mediated induction of PV.1 (Figure 2A and 2B). In addition, co-injection of PV.1 MOs with Xbra also increases the expression of early and late neural marker genes, including *FoxD5a, Ngnr, N-CAM* and *Otx2* and also decreases the PV.1 expression (data not shown). These results significantly evidenced that Xbra catalyzes the inhibition of early neurogenesis in a PV.1-dependent manner. Moreover, these results suggested that PV.1 may be a direct target of Xbra and PV.1 promoter region may contain positive Xbra response elements (XbRE) to facilitate the PV.1-dependent neurogenesis inhibition.

### Identification of Xbra response element (XbRE) within the PV.1 promoter

To identify whether PV.1 promoter region contains cis-acting XbRE elements within 5’-flanking region of PV.1 and positively regulates PV.1 transcription activation, we found that Xbra induces the PV.1 transcription while DN-Xbra reduces the PV.1 transcription (Figure 3A and 3B). These results provide evidence that PV.1 promoter region contains putative cis-acting XbRE and positively regulates PV.1 transcription activation to inhibit the early neurogenesis. In reporter assay of different serially deleted promoter construct found that Xbra increases the relative promoter activity of different serially deleted promoter constructs of PV.1 up to 1.5-4-fold (Figure 3C-3E). These results strongly evidence that PV.1 (−103) promoter construct contains cis-acting XbRE to trigger the PV.1 transcriptional activation and inhibits the early neurogenesis.

A study documented the conserved cis-acting binding response elements (TCACACCT) for the T-box domain containing transcription factor in *Drosophila* (Conlon et al., 2001). In addition, recent study observed that Xbra modulates the expression of its target genes in *Drosophila* embryonic cells in a dose-dependent manner through binding within a consensus sequence (A/G)(A/T)(A/T)NTN(A/G)CAC(C/T)T (Kusch et al., 2002). We further mapped out the cis-acting XbRE within PV.1 (−103) promoter constructs and observed that PV.1 (−103) promoter construct contains a putative cis-acting XbRE (ATCACACTT, within −70bp~-62bp) upstream of putative transcription initiation site (Figure 3F). To test whether putative XbRE positively regulates the PV.1 transcription activation, the loss-of-function study resulted that putative XbRE positively regulates the PV.1 transcription activation (Figure 3G and 3F). These results collectively provide evidence that PV.1 (−103) promoter construct contains cis-acting XbRE which actively participates in transcriptional activation of PV.1 and inhibits the early neurogenesis. Thus, we assumed that whether Xbra directly binds within PV.1 promoter region to activate PV.1 transcription. The ChIP assay analysis confirmed that Xbra directly binds within the proximal region of the endogenous PV.1 promoter (Figure 3H and I). These results collectively evidenced that PV.1 promoter comprises cis-acting XbRE which positively regulates the PV.1 transcription and Xbra directly binds within the proximal region of the PV.1 promoter to inhibit the early neurogenesis.

### Xbra and smad-1 synergistically regulate the PV.1 transcription activation

Our previous study documented that PV.1 promoter region contains the cis-acting BRE (CAGACT, −180bp to −162bp) within 5’-flanking region of putative transcription initiation site of PV.1 (Lee et al., 2011). Moreover, PV.1 is a direct target of BMP signaling and positively regulates the transcriptional activation of PV.1 to inhibit the early neurogenesis (Lee et al., 2011). Another study documented that C-terminal mutated Smad-1 at S378N, Y336D and Y343D abolishes the interaction of Smad-1 with Xbra and C-terminal phosphorylated Smad-1 plays a significant role in Smad-1-Xbra interaction (Messenger et al., 2005). However, the regulatory function of Smad-1-Xbra interaction was unknown. Messenger et al., 2005 suggested that Smad-1-Xbra interaction may induce the expression of the ventral genes (Messenger et al., 2005). Our study reported that BMP-4/Smad-1 and FGF/Xbra signaling cis-acting response elements, BRE and XbRE presents in close vicinity within the proximal region of the PV.1 promoter. Using this hypothesis, we assumed that Xbra and Smad-1 mediates a signaling crosstalk between BMP-4/Smad-1 and FGF/Xbra signaling in context to PV.1 transcription activation. Our study provides evidence that Xbra and Smad-1 strongly increase the promoter activity of PV.1 (−180) promoter construct, which contains both BRE and XbRE, and cooperates synergistically to activate the PV.1 transcription for inhibiting early neurogenesis (Figure 4A and 4B). We further hypothesized that whether XbRE plays a role in the synergistic regulation of PV.1 transcription. The loss-of-function study of XbRE resulted that Xbra and Smad-1-mediated synergistic activation of PV.1 (−103) promoter construct is completely abolished with PV.1 (−103)mXbRE (Figure 4C). This finding suggested that XbRE plays a significant role in a synergistic activation of PV.1 transcription.

We further raised questions that whether XbRE or BRE, which one plays a more crucially role in synergistic activation of PV.1 transcription to inhibit the neurogenesis. Thus, our study evidenced that relative promoter activity of BRE mutated PV.1 (−180)mBRE (up to 2.5-fold) declines more remarkably in Xbra-Smad-1-injected embryos as compared to PV.1(−180)mXbRE (up to 8.5-fold) (Figure 4D and 4E). These results concluded that BRE plays a more significant role in a synergistic regulation of PV.1 transcription activation as compared to XbRE. Moreover, these finding collectively suggested that BRE-mediated transcriptional activation of PV.1 inhibits early neurogenesis more efficiently and plays a crucial role in signaling crosstalk of BMP-4/Smad-1 and FGF/Xbra during *Xenopus* embryos. Moreover, the doubly loss-of-function study in PV.1 (−180) promoter construct showed that Smad-1 and Xbra-mediated synergistic activation of PV.1 transcription are completely abolished with PV.1 (−180)m(BRE+XbRE) promoter construct (Figure 4F). These results concluded that both XbRE and BRE cis-acting response elements cooperate synergistically to regulate the PV.1 transcription activation and inhibit the early neurogenesis in the ectoderm. In addition, BRE plays a more profound role in signaling crosstalk, initiated by BMP-4/Smad-1 and FGF/Xbra in context to transcriptional activation of PV.1. Additionally, PV.1 may be a novel target of Smad-1 and Xbra interaction in a Vent family because the physical interaction of Smad-1 and Xbra does not regulate the Xvent2 expression in a synergistic manner (S1).

A study documented that C-terminal phosphorylation of Smad-1 plays a significant role in the physical interaction of Smad-1 and Xbra. Smad-1 physically interacts with an N-terminal domain of Xbra while C-terminal phosphorylated mutant Smad-1 (S378N, Y336D, and Y343D) does not interact with Xbra (Messenger et al., 2005). Thus, we assumed that whether Xbra and Smad-1 interact with each other to establish the signaling crosstalk for synergistic regulation of PV.1 transcription activation. The immunoprecipitation assay showed that Xbra physically interacts with C-terminal phosphorylated Smad-1 and phospho-mimic 3SD (S462D, S463D, and S465D) Smad-1 while Xbra interaction with phospho-dead 3SA (S462A, S463A, and S465A) Smad-1 is not reported (Figure 4G and 4H). This study concludes that Smad-1 C-terminal phosphorylation plays a significant role in Xbra and Smad-1 interaction and establishes a signaling crosstalk between BMP-4/Smad-1 and FGF/Xbra to activate the PV.1 transcription in a synergistic manner for inhibiting the early neurogenesis in the ectoderm.

### Physical binding of Smad-1 and Xbra stimulates promoter activity of PV.1 and is dependent on DNA binding, but not absolutely require DNA binding for cooperative stimulation

Above collective findings showed the evidence that Xbra and Smad-1 cooperate synergistically to regulate the transcriptional activation of PV.1 and establishes a signaling crosstalk between BMP-4/Smad-1 and FGF/Xbra in context to activation of PV.1 transcription. Thus, we hypothesized that whether Smad-1 and Xbra increase the DNA binding of their respective interacting partners with the endogenous PV. 1 promoter region for synergistic regulation of PV. 1 transcription. Moreover, whether DNA binding of Xbra and Smad-1 is absolutely required for synergistic activation of PV.1 transcription. In ChIP assay found that Smad-1 increases Xbra binding with the endogenous promoter region of PV.1 (Figure 5A and B). Moreover, we observed that Xbra could also increase Smad-1 binding with the endogenous promoter region of PV.1 (Figure 5C and D). These results suggested that Smad-1 and Xbra not only triggers the promoter activity of PV.1 transcription in a synergistic manner but also promotes the DNA binding of their interacting partner and facilitates a signaling crosstalk in context to PV.1 transcriptional activation. Surprisingly, we observed that Smad-1 and Xbra binding to the endogenous PV.1 promoter are only required for transcriptional activation of PV.1, but this binding is not absolutely required for cooperative stimulation of PV.1 transcriptional activation. Taken together, all collective findings suggested that the 5’-flanking region of putative transcription start site of PV.1 contains cis-acting XbRE within −70bp~-62bp upstream of putative transcription initiation site of PV.1. Furthermore, we found that Xbra is a positive regulator of PV.1 transcription and inhibits the early neurogenesis in a PV.1-dependent manner.

Moreover, Xbra and Smad-1 cooperates synergistically to regulate the PV.1 transcription activation positively and inhibits the early neurogenesis in ectoderm explant. In addition, BRE plays a more significant role in synergistic regulation the PV.1 transcription rather than XbRE and inhibits the early neurogenesis more efficiently. We also observed that FGF/Xbra and BMP-4/Smad-1 establishes a signaling crosstalk to inhibit the early neurogenesis. Additionally, Xbra and Smad-1 directly binds within PV.1 promoter region and enhances the binding of their respective interacting partner with the endogenous PV.1 promoter, but this promoter DNA binding is not absolutely required for synergistic regulation of PV.1 transcription activation during *Xenopus* development.

In last, we proposed a putative systematic model for Xbra-Smad-1-mediated synergistic regulation of PV.1 (−180) promoter constructs and its transcriptional activation. Moreover, BMP-4/Smad-1 and FGF/Xbra-mediated signaling crosstalk in context to PV.1 transcription activation and in neurogenesis inhibition (Figure 6).

**Figure 6:**
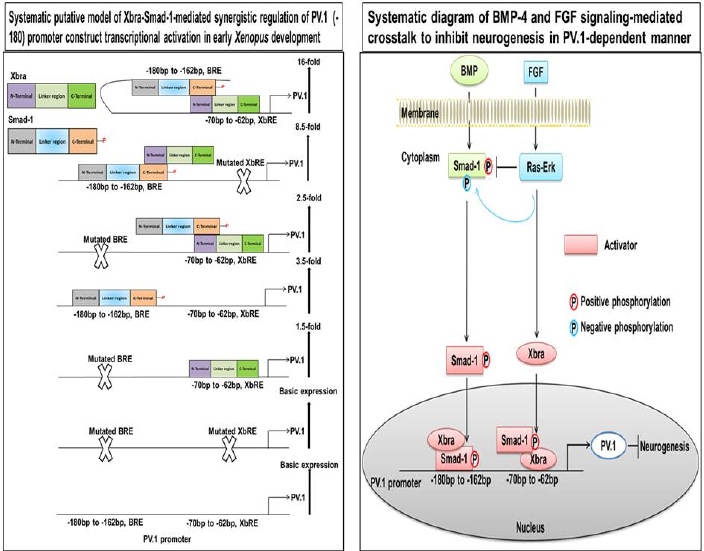
The systematic putative model of Xbra-Smad-1-mediated synergistic regulation of PV.1 transcriptional activation and neurogenesis inhibition during early *Xenopus*development. (A) Smad-1 and Xbra-mediated putative transcriptional regulation of PV1 (−180) promoter construct is shown in a synergistic manner. BMP-4/Smad-1 and FGF/Xbra-mediated signaling crosstalk in context to PV1 transcription activation to inhibit the early neurogenesis during *Xenopus* development.

## Material and methods

### Ethics Statement

Institutional Animal Care and Use Committee (IACUC) approval is not required for the experimental use of amphibians or reptiles in Korea. All members of our research group are attended the both educational and training courses to appropriate care and usage of experimental animals. Adult *X. laevis* were grown in 12 hrs light/dark (LD 12:12 h) cycles at 18^0^C according to the guidelines of Institutes of Laboratory Animal Resources that works for laboratory animal maintenance.

### DNA and RNA preparation

cDNAs encoding Dominant-negative BMP receptor (DNBR), Smad-1 (WT): SP6, Asp718, and its mutants Smad-1 (3SD): SP6, Asp718, and Smad-1 (3SA): SP6, Asp718 were all subcloned into the pSP64T expression vector while Xbra: SP6, Asp718 was subcloned into the pCS2+ expression vector (Lee et al., 2011). Each vector was linearized with the appropriate restriction enzyme and used for in-vitro transcription using the MEGAscript kit according to manufacturer’s instructions (Ambion, Austin, TX). Synthetic mRNAs were quantified by spectrophotometer at 260/280nm (SPECTRA max, Molecular Devices).

### Cloning of PV.1A genomic DNA

The cloning of PV.1A genomic DNA (gDNA) was performed into the pBluescript SK(-) plasmid (Stratagene, Cedar Creek, TX) as described by Lee et al., 2011 (Lee et al., 2011).

### PV.1 promoter constructs

The 2.5 kb of 5’-flanking region of positive clone was subcloned into the pGL-2 basic plasmid (Promega, Madison, WI) and was designated the −2525bp construct. Serially deleted PV.1 promoter mutants and triple-repeat BMP-4-response element (BRE) were generated from −2525bp construct and subcloned into a pGL-2 basic plasmid by PCR amplification (Table 1) according to Lee et al. 2011 (Lee et al., 2011).

**Table 1.**
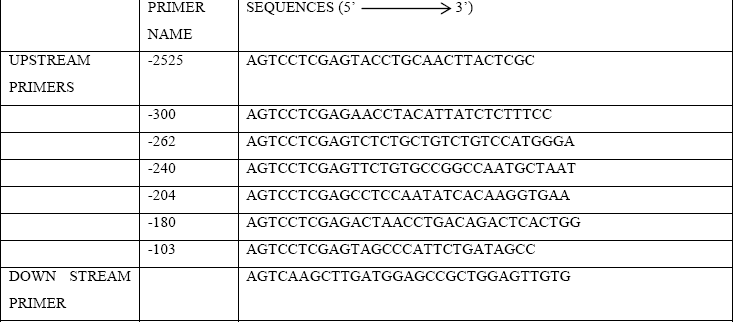
Primers used for serially-deleted reporter gene constructs.

### Embryo injection and explants culture

*Xenopus* embryos were injected after in vitro fertilization of eggs that induced by injection of 500 units of human chorionic gonadotropin (Sigma, St. Louis, MO). RNAs were injected into the animal pole at 1-cell stage embryos and harvested into 30% MMR. Then, animal cap (AC) dissected from injected embryos at stage 8.0-8.5 and incubated in 1X L-15 growth medium (Gibco) till stage 11 for RT-PCR.

### RNA isolation and RT-PCR

Xbra mRNA (1 ng) was injected into the animal pole at 1-cell stage of *Xenopus* embryos and harvested into 30% MMR solution with respect to control non-injected embryos till stage 8. Animal caps were then dissected from the injected and non-injected embryos and incubated until stage 11 or 24 into 1X L-15 growth medium. Total RNA was isolated from whole embryos or animal caps using RNA-bee reagent following the manufacturer’s instructions (TEL-TEST, Friendwood, Texas) and treated with DNase I remove gDNA contamination. RT-PCR was performed with Superscript II (Invitrogen, Carlsbad, CA), as described by the manufacturer, with 2 mg total RNA per reaction. PCR was performed according to the following conditions: 30 seconds at 94^0^C, 30 seconds at each annealing temperature, 30 seconds at 72^0^C; 20-28 cycles of amplification (Table 2).

**Table 2.**
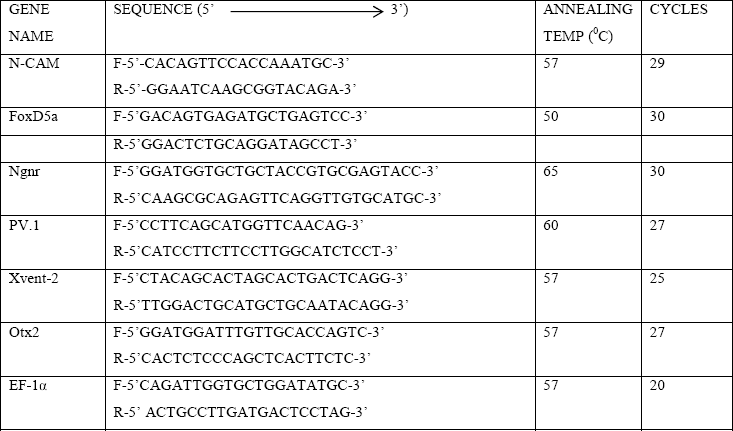
Primers used for RT-PCR amplification.

### Luciferase assays

Relative luciferase reporter gene activities were measured using luciferase assay system according to manufacturer’s instructions (Promega, Madison, WI). Five different groups of embryos (3 embryos per group) were harvested and homogenized in 10μl lysis buffer per animal cap. 10μL embryos homogenate were assayed with 40μL luciferase substrate and determined the reporter gene activity by the luminometer (EG & G Berthold, Bad Wildbad, Germany). All experiments were repeated at least three times using independently derived sample sets.

### Site-directed mutagenesis

Mutagenesis was performed by a site-directed mutagenesis (Muta-Direct™ iNtRON Biotechnology) kit using the several oligonucleotides in accordance with instructions (Table 3).

**Table 3.**
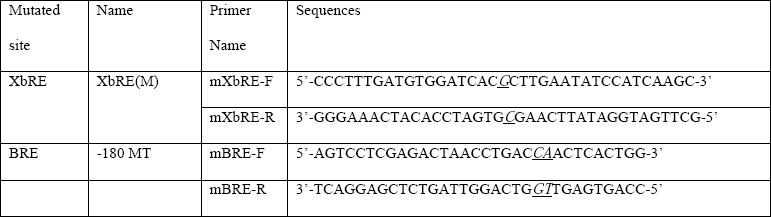
Primers used for site-direct mutagenesis gene constructs.

### Immunoprecipitation

Embryos were co-injected with Xbra-myc mRNA at one cell stage with Smad-1-Flag, 3SD, and 3SA in three different groups and injected embryos were collected at stage 11. They were then homogenized in lysis IP buffer (1M Tris [7.4]), 150mM NaCl, 1% NP-40, 10% Triton-X, 0.5M EDTA, 50% glycerol, 50mM NaF, 1mM Na3VO4, 15mM Glycerophosphate and 200X phosphatase inhibitors (PMSF [Sigma, cat-7626], Pepstatin A [Sigma, cat-P4265], Leupeptin [Sigma, cat-L0649], Benzamidine [Sigma, cat-B6506]. Cell lysates were cleared by centrifugation and cleared lysate incubated with the C-Myc polyclonal antibody (Santa Cruz Biotechnology, SC-789) for overnight at 4^0^C, immunocomplexes were precipitated by using protein A/G beads plus (Santa Cruz Biotechnology, SC-2003). The proper amount of precipitated beads-protein complex was boiled in sample buffer, and resolved by electrophoresis on 10% SDS-polyacrylamide gels. Western blotting of Flag-Smad-1, Flag-Smad-1(3SD), and Flag-Smad-1(3SA) was performed by using an anti-Flag monoclonal antibody (Sigma, F−1804) and secondary antibody anti-mouse purchased by Stressgen (SAB-100). Immune complexes were visualized by using an ECL detection kit (GE healthcare).

### Chromatin immune-precipitation (ChIP)

Chromatin immunoprecipitation assay was performed as described (Blythe et al., 2009). Embryos were injected at the one-cell stage with mRNA encoding Xbra-Myc and Smad-1-Flag (1ng per embryo) either separately or in combination. Injected embryos were collected at stage 11 (100 embryos/sample) and processed according to the protocol. Polyclonal C-Myc (Santa Cruz Biotechnology, SC-789) and anti-Flag monoclonal antibody (Sigma, F−1804) were added to immunoprecipitate the chromatin. Moreover, normal rabbit IgG (Santa Cruz Biotechnology, SC-2027) and normal mouse IgG (Santa Cruz Biotechnology, SC-2025). PCR were performed with immunoprecipitated fragmented chromatin using PV.1 (−180 and −103) promoter region primers. Primers were shown in Table 1 and 2.

### Morpholino Oligos (MOs)

PV.1 morpholino oligos (MOs, Genetools, LLC) is an anti-sense oligodeoyynucletides were used for loss-of-function study. This morpholino was designed against 5’ UTR and/or the start site of transcription initiation. The sequence of PV.1 MOs was as follows:

> PV.1 MOs: 5’- AATCTTTGTTTGAACCATGCTGAAGG -3’

MOs were warmed at 55^0^C for 5 min and keep at 37^0^C until injection the embryos to avoid the clogging of microinjection needles. MOs were injected with 10ng per embryos.

### Nucleotide sequence accession number

The PV.1 (accession number; AF133122) cDNA sequence has been submitted to GenBank.

### Whole mount in-situ hybridization

Embryos were injected with mRNAs at one-cell stage and performed whole-mount in situ hybridization at stage 11 using standard methods with anti-sense probes for Xvent2 (Yoon et al., 2014a).

### Statistical analysis

Data were analyzed by unpaired two-tailed Student’s t test using GraphPad Prism4. Asterisks denote **: p ≤ 0.01, ***: p ≤ 0.001, n.s.: not significant.

### Conflicts of authors

There is no author conflict to disclose.

### Funding

This research was supported by National Research Foundation of Korea (NRF), funded by the Ministry of Education, Science and Technology (NRF-2016R1D1A1B02008770) and (NRF-2016M3A9B8914057).

## References

Ault, K. T., Dirksen, M. L. and Jamrich, M. (1996) ‘A novel homeobox gene PV.1 mediates induction of ventral mesoderm in Xenopus embryos’, Proc Natl Acad Sci U S A 93(13): 6415–20.

Blythe, S. A., Reid, C. D., Kessler, D. S. and Klein, P. S. (2009) ‘Chromatin immunoprecipitation in early Xenopus laevis embryos’, Dev Dyn 238(6): 1422–32.

Bouwmeester, T., Kim, S., Sasai, Y., Lu, B. and De Robertis, E. M. (1996) ‘Cerberus is a head-inducing secreted factor expressed in the anterior endoderm of Spemann’s organizer’, Nature 382(6592): 595–601.

Conlon, F. L., Fairclough, L., Price, B. M., Casey, E. S. and Smith, J. C. (2001) ‘Determinants of T box protein specificity’, Development 128(19): 3749–58.

Dale, L. and Wardle, F. C. (1999) ‘A gradient of BMP activity specifies dorsal-ventral fates in early Xenopus embryos’, Semin Cell Dev Biol 10(3): 319–26.

Delaune, E., Lemaire, P. and Kodjabachian, L. (2005) ‘Neural induction in Xenopus requires early FGF signalling in addition to BMP inhibition’, Development 132(2): 299–310.

Fainsod, A., Deissler, K., Yelin, R., Marom, K., Epstein, M., Pillemer, G., Steinbeisser, H. and Blum, M. (1997) ‘The dorsalizing and neural inducing gene follistatin is an antagonist of BMP-4’, Mech Dev 63(1): 39–50.

Gamse, J. and Sive, H. (2000) ‘Vertebrate anteroposterior patterning: the Xenopus neurectoderm as a paradigm’, Bioessays 22(11): 976–86.

Gawantka, V., Delius, H., Hirschfeld, K., Blumenstock, C. and Niehrs, C. (1995) ‘Antagonizing the Spemann organizer: role of the homeobox gene Xvent-1’, EMBO J 14(24): 6268–79.

Hawley, S. H., Wunnenberg-Stapleton, K., Hashimoto, C., Laurent, M. N., Watabe, T., Blumberg, B. W. and Cho, K. W. (1995) ‘Disruption of BMP signals in embryonic Xenopus ectoderm leads to direct neural induction’, Genes Dev 9(23): 2923–35.

Hemmati-Brivanlou, A. and Melton, D. (1997) ‘Vertebrate neural induction’, Annu Rev Neurosci 20: 43–60.

Hemmati-Brivanlou, A. and Thomsen, G. H. (1995) ‘Ventral mesodermal patterning in Xenopus embryos: expression patterns and activities of BMP-2 and BMP-4’, Dev Genet 17(1): 78–89.

Hwang, Y. S., Lee, H. S., Roh, D. H., Cha, S., Lee, S. Y., Seo, J. J., Kim, J. and Park, M. J. (2003) ‘Active repression of organizer genes by C-terminal domain of PV.1’, Biochem Biophys Res Commun 308(1): 79–86.

Hwang, Y. S., Seo, J. J., Cha, S. W., Lee, H. S., Lee, S. Y., Roh, D. H., Kung Hf, H. F., Kim, J. and Ja Park, M. (2002) ‘Antimorphic PV.1 causes secondary axis by inducing ectopic organizer’, Biochem Biophys Res Commun 292(4): 1081–6.

Kim, J., Lin, J. J., Xu, R. H. and Kung, H. F. (1998) ‘Mesoderm induction by heterodimeric AP-1 (c-Jun and c-Fos) and its involvement in mesoderm formation through the embryonic fibroblast growth factor/Xbra autocatalytic loop during the early development of Xenopus embryos’, J Biol Chem 273(3): 1542–50.

Kusch, T., Storck, T., Walldorf, U. and Reuter, R. (2002) ‘Brachyury proteins regulate target genes through modular binding sites in a cooperative fashion’, Genes Dev 16(4): 518–29.

Ladher, R., Mohun, T. J., Smith, J. C. and Snape, A. M. (1996) ‘Xom: a Xenopus homeobox gene that mediates the early effects of BMP-4’, Development 122(8): 2385–94.

Lee, H. S., Lee, S. Y., Lee, H., Hwang, Y. S., Cha, S. W., Park, S., Lee, J. Y., Park, J. B., Kim, S., Park, M. J. et al. (2011) ‘Direct response elements of BMP within the PV.1A promoter are essential for its transcriptional regulation during early Xenopus development’, PLoS One 6(8): e22621.

Lee, H. S., Park, M. J., Lee, S. Y., Hwang, Y. S., Lee, H., Roh, D. H., Kim, J. I., Park, J. B., Lee, J. Y., Kung, H. F. et al. (2002) ‘Transcriptional regulation of Xbr-1a/Xvent-2 homeobox gene: analysis of its promoter region’, Biochem Biophys Res Commun 298(5): 815–23.

Melby, A. E., Clements, W. K. and Kimelman, D. (1999) ‘Regulation of dorsal gene expression in Xenopus by the ventralizing homeodomain gene Vox’, Dev Biol 211(2): 293–305.

Messenger, N. J., Kabitschke, C., Andrews, R., Grimmer, D., Nunez Miguel, R., Blundell, T. L., Smith, J. C. and Wardle, F. C. (2005) ‘Functional specificity of the Xenopus T-domain protein Brachyury is conferred by its ability to interact with Smad1’, Dev Cell 8(4): 599–610.

Onichtchouk, D., Gawantka, V., Dosch, R., Delius, H., Hirschfeld, K., Blumenstock, C. and Niehrs, C. (1996) ‘The Xvent-2 homeobox gene is part of the BMP-4 signalling pathway controlling [correction of controling] dorsoventral patterning of Xenopus mesoderm’, Development 122(10): 3045–53.

Papalopulu, N. and Kintner, C. (1996) ‘A Xenopus gene, Xbr-1, defines a novel class of homeobox genes and is expressed in the dorsal ciliary margin of the eye’, Dev Biol 174(1): 104–14.

Pera, E. M., Ikeda, A., Eivers, E. and De Robertis, E. M. (2003) ‘Integration of IGF, FGF, and anti-BMP signals via Smad1 phosphorylation in neural induction’, Genes Dev 17(24): 3023–8.

Piccolo, S., Sasai, Y., Lu, B. and De Robertis, E. M. (1996) ‘Dorsoventral patterning in Xenopus: inhibition of ventral signals by direct binding of chordin to BMP-4’, Cell 86(4): 589–98.

Rao, Y. (1994) ‘Conversion of a mesodermalizing molecule, the Xenopus Brachyury gene, into a neuralizing factor’, Genes Dev 8(8): 939–47.

Rastegar, S., Friedle, H., Frommer, G. and Knochel, W. (1999) ‘Transcriptional regulation of Xvent homeobox genes’, Mech Dev 81(1-2): 139–49.

Saka, Y., Tada, M. and Smith, J. C. (2000) ‘A screen for targets of the Xenopus T-box gene Xbra’, Mech Dev 93(1-2): 27–39.

Sasai, Y., Lu, B., Steinbeisser, H. and De Robertis, E. M. (1995) ‘Regulation of neural induction by the Chd and Bmp-4 antagonistic patterning signals in Xenopus’, Nature 376(6538): 333–6.

Schier, A. F. (2001) ‘Axis formation and patterning in zebrafish’, Curr Opin Genet Dev 11(4): 393–404.

Schmidt, J. E., von Dassow, G. and Kimelman, D. (1996) ‘Regulation of dorsal-ventral patterning: the ventralizing effects of the novel Xenopus homeobox gene Vox’, Development 122(6): 1711–21.

Trindade, M., Tada, M. and Smith, J. C. (1999) ‘DNA-binding specificity and embryological function of Xom (Xvent-2)’, Dev Biol 216(2): 442–56.

Weisinger, K., Wilkinson, D. G. and Sela-Donenfeld, D. (2008) ‘Inhibition of BMPs by follistatin is required for FGF3 expression and segmental patterning of the hindbrain’, Dev Biol 324(2): 213–25.

Wilson, P. A. and Hemmati-Brivanlou, A. (1995) ‘Induction of epidermis and inhibition of neural fate by Bmp-4’, Nature 376(6538): 331–3.

Wilson, S. I., Rydstrom, A., Trimborn, T., Willert, K., Nusse, R., Jessell, T. M. and Edlund, T. (2001) ‘The status of Wnt signalling regulates neural and epidermal fates in the chick embryo’, Nature 411(6835): 325–30.

Xu, R. H., Kim, J., Taira, M., Zhan, S., Sredni, D. and Kung, H. F. (1995) ‘A dominant negative bone morphogenetic protein 4 receptor causes neuralization in Xenopus ectoderm’, Biochem Biophys Res Commun 212(1): 212–9.

Yoon, J., Kim, J. H., Kim, S. C., Park, J. B., Lee, J. Y. and Kim, J. (2014a) ‘PV.1 suppresses the expression of FoxD5b during neural induction in Xenopus embryos’, Mol Cells 37(3): 220–5.

Yoon, J., Kim, J. H., Lee, S. Y., Kim, S., Park, J. B., Lee, J. Y. and Kim, J. (2014b) ‘PV.1 induced by FGF-Xbra functions as a repressor of neurogenesis in Xenopus embryos’, BMB Rep.

Zimmerman, L. B., De Jesus-Escobar, J. M. and Harland, R. M. (1996) ‘The Spemann organizer signal noggin binds and inactivates bone morphogenetic protein 4’, Cell 86(4): 599–606.

